# Gene flow throughout the evolutionary history of a polymorphic and generalist clownfish

**DOI:** 10.1101/2023.10.19.563036

**Authors:** Sarah Schmid, Marianne Bachmann Salvy, Alberto Garcia Jimenez, Joris A. M. Bertrand, Fabio Cortesi, Sara Heim, Filip Huyghe, Glenn Litsios, Anna Marcionetti, James O’Donnell, Cynthia Riginos, Valerio Tettamanti, Nicolas Salamin

## Abstract

Even seemingly homogeneous on the surface, the oceans display high environmental heterogeneity across space and time. Indeed, different soft barriers structure the marine environment, which offers an appealing opportunity to study various evolutionary processes such as population differentiation and speciation. Here, we focus on *Amphiprion clarkii* (Actinopterygii; Perciformes), the most widespread of clownfishes that exhibits the highest colour polymorphism. Clownfishes can only disperse during a short pelagic larval phase before their sessile adult lifestyle, which might limit connectivity among populations, thus facilitating speciation events. Consequently, the taxonomic status of *A. clarkii* has been under debate. We used whole-genome resequencing data of 67 *A. clarkii* specimens spread across the Indian and Pacific Oceans to characterise the species’ population structure, demographic history, and colour polymorphism. We found that *A. clarkii* spread from the Indo-Pacific Ocean to the Pacific and Indian Oceans following a stepping-stone dispersal and that gene flow was pervasive throughout its demographic history. Moreover, edge populations exhibited more similar colouration patterns compared to central populations. However, we demonstrate that colour polymorphism is not associated with population structure, thus, colour phenotype is unreliable in assessing the taxonomic status of *A. clarkii*. Our study further highlights the power of whole-genome comparative studies to determine the taxonomy of geographically wide-ranging and phenotypically diverse species, supporting the status of *A. clarkii* as a single species.

## INTRODUCTION

Marine ecosystems are usually thought to be highly connected (Conover et al., 2006; Shanks et al., 2003) due to the apparent lack of barriers to gene flow and the large population sizes of many marine species (Palumbi, 1992; Waples, 1998). Despite this seeming absence of boundaries, oceans and seas constitute structured ecosystems. Past geographical events, water salinity and temperature, as well as oceanic currents, are some of the multiple environmental factors that might form partial or “soft” barriers in the marine environment (Belanger et al., 2012; Choo et al., 2021; Marko, 2004; Meyer et al., 2005; Oleksiak, 2019; White et al., 2010). Studying population connectivity in this environment is appealing since it promises to provide critical insights into the impact of soft barriers on population structure and speciation.

The Indo-Pacific region is widely recognized as the centre of marine biodiversity, with the Coral Triangle being a significant biodiversity hotspot (Hughes et al., 2002, Renema et al., 2008; Lohman et al., 2011) not only for fishes, but also for corals and other invertebrate species (Hoeksema, 2007; Allen, 2008). However, the factors that have led to this consistent biogeographic pattern still need to be fully understood (Pellissier et al., 2014). One approach to understanding the history of this region is to study intraspecific variation to make inferences about the factors that may have shaped the distribution of marine species. This can shed light on questions such as connectivity within a species, how much they have spread, their geographical origins, and whether edge locations are peripheral in terms of genetic diversity (Godhe and Rynearson, 2017; Buckley et al., 2022; Arroyo-Correa et al., 2023). Nevertheless, obtaining sufficient population data in marine environments to infer the processes that lead to population divergence and structure remains challenging (Keyse et al., 2014). Consequently, the number of marine studies is limited compared to their terrestrial counterparts. Early investigations into the population structure of marine animals revealed that they tend to show limited genetic structure (Conover et al., 2006; Waples, 1998). However, most of these studies were based on a few genetic markers. More recently, the transition to whole-genome analyses enabled the detection of previously unidentified population differentiation (Allendorf et al., 2010; Ellegren, 2014). It also allowed researchers to go beyond the description of these patterns to test hypotheses about the processes leading to population structure (Hellberg, 2009). Such extensive genomic coverage further enhanced the estimation of population-genetic parameters, such as migration rates or effective population sizes. It enabled the formulation of neutral expectations with more confidence and improved inferences about population demography (Allendorf et al., 2010), considerably changing the perspective on marine ecosystems. Hence, in recent years there has been a switch from a vision of well-mixed marine populations to a view of populations with restricted gene flow displaying considerable levels of differentiation and intraspecific variability (Barabás and D’Andrea, 2016; Messer et al., 2016; Rudman et al., 2015; van der Ven et al., 2021; Wood and Brodie, 2016). This change of paradigm challenges past assumptions about marine species dispersal and calls for an improved understanding of the factors influencing gene flow and their resulting evolutionary implications.

In this context, clownfishes – also known as anemonefishes (Actinopterygii; Perciformes; Amphiprioninae) – represent an exciting group of reef fishes for population genomic studies due to their specific life-history traits. In the wild, clownfishes have formed mutualistic relationships with ten possible sea anemone species, with specialist species that only interact with a subset of anemones and general species that may access the full range of hosts (Litsios et al. 2012). Moreover, clownfishes dispersal is restricted by their short pelagic larval duration (PLD), which lasts from 10 to 15 days (Fautin et al., 1997; Roux et al., 2019; Roux et al., 2020), and exhibit high levels of self-recruitment (Beldade et al., 2012; Buston et al., 2012; Jones et al., 2005; Madduppa et al., 2014). This pelagic phase results in a general dispersal range of around five to one hundred kilometres (Salinas-de-León, Jones, and Bell, 2012), which is much shorter compared to most other damselfish species (Thresher, Colin, and Bell, 1989). It needs to be noted that long-distance dispersal has been recorded in some clownfish species, presumably due to strong oceanic currents (e.g., over 400 km in *Amphiprion omanensis*; Simpson et al., 2014). Besides a low dispersal ability and strong habitat specificity, clownfishes are monogamous protandrous species whose social structure is controlled by a strict hierarchy. These strong constraints put into question the demographic history and the microevolutionary processes involved in the range expansions observed in clownfishes (Huyghe and Kochzius, 2017; Litsios et al., 2014), with some species displaying an unexpectedly broad geographic range.

*Amphiprion clarkii* (Bennett, 1830) - a highly polymorphic and generalist clownfish species - exhibits the broadest geographic range among all clownfishes (Fautin and Allen, 2007). Its range extends from the Micronesian islands of the Pacific Ocean to the Persian Gulf and stretches longitudinally from the Japanese coast to the Great Barrier Reef (Fautin et al., 1997). Nevertheless, the geographical origin of the species, the potential impact of sea surface currents and geographical barriers on the connectivity between populations, and the evolutionary processes behind its wide distribution remain unclear (Litsios et al., 2014). Moreover, *A. clarkii* is considered an “extreme generalist” because of its ability to interact with all ten sea anemone hosts (Ollerton et al., 2007). This characteristic could either facilitate connectivity among populations due to the increased host anemone availability or – on the opposite – might reduce gene flow among neighbouring populations due to habitat-specific host selection. Finally, this species displays a high level of colour polymorphism, which has led to considerable confusion regarding their identification. Indeed, the background colour can range from light orange to entirely dark, and the number of emblematic white bars varies among individuals (Bell et al., 1982; Fautin et al., 1997). All these characteristics started a several decade-long debate about the potential for *A. clarkii* to consist of a complex of multiple species. Phylogenetic reconstructions based on a few genetic markers showed contradictory results at the molecular level. A. *clarkii* specimens from the Indo-Pacific Ocean clustered by colouration rather than geographic origin (Litsios et al., 2014; Rolland et al., 2018) while colour divergent *A. clarkii* morphotypes from the Indian Ocean did not differ significantly at the molecular level (Devi et al., 2021). Moreover, the evolutionary processes involved in the remarkable colour variation, geographical range, and mutualistic interactions displayed by *A. clarkii* remain to be deciphered to provide certainty about the taxonomic status of the species.

In this study, we analysed the most extensive sampling of *A. clarkii* populations to date using whole-genome resequencing approaches to assess the taxonomic status of *A. clarkii* and thus evaluate the impact of soft barriers on the species. We inferred the demographic history and population structure and computed genome-wide phenotype associations to test for demographic expansions, restricted gene flow, and the genetic association of colour polymorphism in *A. clarkii* populations. The broad geographical range of *A. clarkii* raises questions about the geographic origin of the species and the dynamic of the range expansion that followed. We tested two different hypotheses that propose either an origin in the Indo-Pacific Ocean – following the ‘Center of Origin’ hypothesis (Briggs, 2003; Gaboriau *et al*., 2017) – or in the Papua New Guinea-Solomon Islands area, which could lead to a subsequent west-ward expansion through a stepping-stone process. We further hypothesised that the unique characteristics of *A. clarkii* compared to other clownfishes may indicate the presence of cryptic genetic differentiation, questioning its status as a single species. We thus estimated gene flow among populations to determine whether gene flow occurred throughout the evolutionary history of *A. clarkii*, was restricted to a brief period following population splits, occurred after secondary contact, or occurred recently. Additionally, we investigated the presence of genomic regions displaying a signature of local adaptation. Finally, observed colour polymorphism could be the cause or consequence of reproductive isolation, leading to an association between potential genetic differences and colour morphs. We conducted genome-wide analyses to test if the genetic structure was associated with specific colour morphs and if genomic differentiation involved loci known to influence colour variation in fishes. Our study demonstrates the potential of whole-genome data to improve our understanding of the evolution of geographically wide-ranging and phenotypically diverse species.

## MATERIAL AND METHODS

### Sampling, library preparation and sequencing

Tissue samples consisting of pieces of dorsal fins of about 1 cm long (fin clips) from 67 *Amphiprion clarkii* specimens and 22 *A. akindynos* (used as outgroup) were collected by various collaborators between 2013 and 2018 from 8 different populations across the Indo-Pacific (Figure 1A; Supplementary Table S1; Appendix 1 for sampling permits) and were stored in ethanol at −20°C. Pictures for morphometry and colour pattern analysis were taken for individuals coming from New Caledonia, Maldives and Indonesia. We extracted the genomic DNA of each fin clip following the DNeasy Blood and Tissue kit standard procedure and performed the final elution twice in 100 µl of AE buffer (QIAGEN, Hombrechtikon, Switzerland). We quantified the extracted DNA using Qubit® 2.0 Fluorometer (Thermo Fisher Scientific, Waltham, USA) and evaluated the integrity by electrophoresis. We followed the TruSeq Nano DNA library prep standard protocol to generate libraries with a 350 base pair insert size for whole-genome paired-end sequencing (Illumina, San Diego, USA). We validated the fragment length distribution of the libraries with a Bioanalyzer (Agilent Technologies, Santa Clara, USA). Sequencing was performed by the Genomic Technologies Facility at the University of Lausanne, Switzerland. Libraries generated in 2017 and 2018 were sequenced on five Illumina HiSeq 2500, 100 paired-end lanes, while libraries prepared in 2019 were sequenced on nine Illumina 4000 HiSeq, 150 paired-end lanes. We aimed for each sample to reach an approximate 10X coverage with both types of libraries.

**Figure 1.**
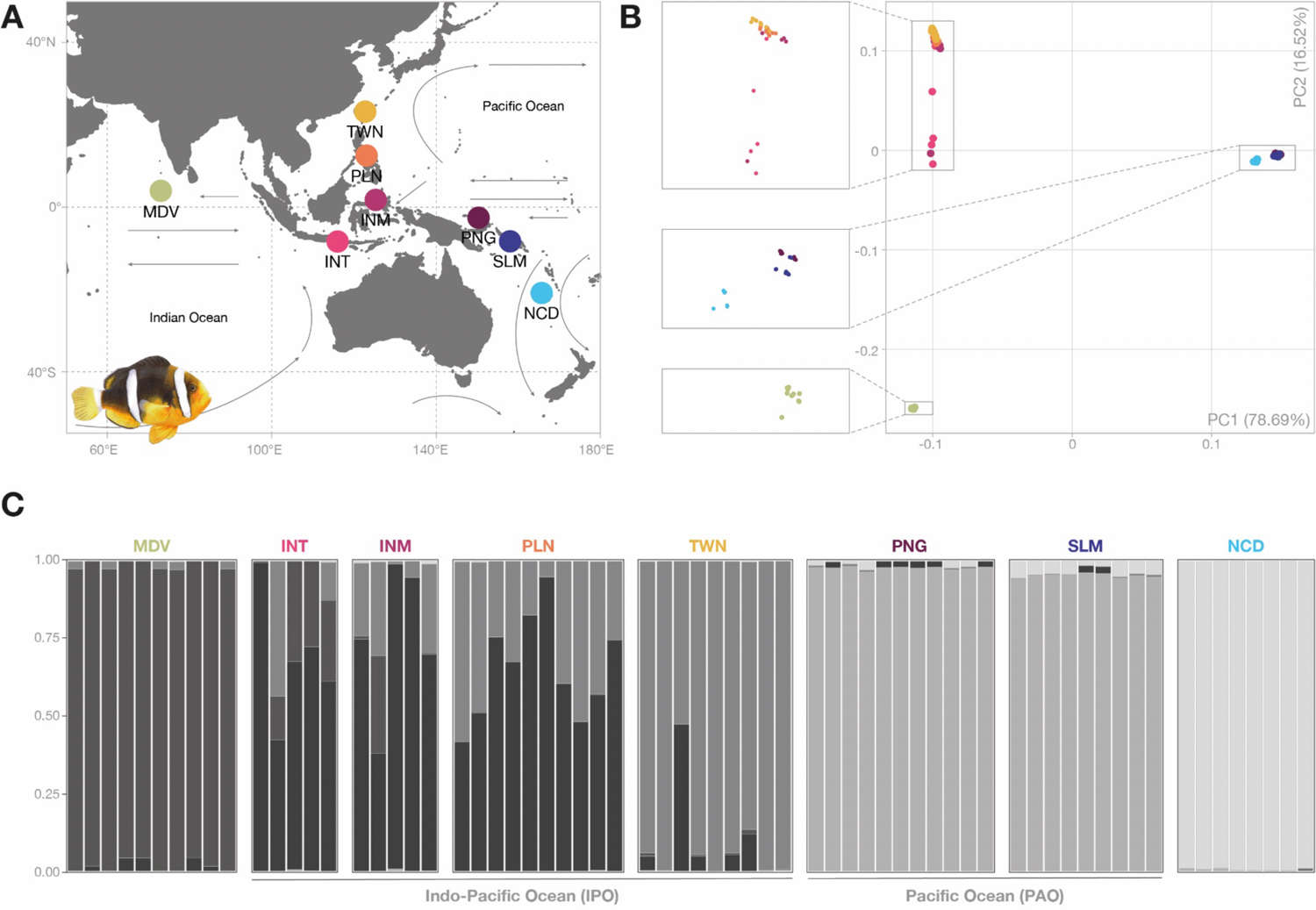
Sampling and population genetic structure of *Amphiprion clarkii*. (A) Coloured circles represent sampling sites for the study: MDV (Maldives), INT (Indonesia, Tulamben), INM (Indonesia, Manado), PLN (Philippines), TWN (Taiwan), PNG (Papua New Guinea), SLM (Solomon Islands), NCD (New Caledonia). Arrows represent major ocean surface currents. (B) Principal component analysis showing a separation of the eastern Pacific populations (PNG, SLM, NCD) from the Indo-Pacific populations (MDV, INM, INT, PLN, TWN) on the first principal component axis (PC1), and further differentiation of the Maldives population (MDV) on the second axis (PC2). (C) Admixture bar plot based on PCANGSD *q*-values for *K*=5. Shades of grey represent the 5 clusters membership for each individual.

### Sequenced data processing, mapping, and genotyping

We trimmed the generated reads to remove adapter sequences using Cutadapt V2.3 (Martin, 2011). We removed reads shorter than 40 bp and with a Phred quality score below 40 with Sickle V1.33 (Joshi and Fass, 2011). We assessed read quality before and after processing with FastQC V0.11.7 (Andrews, 2010). We mapped the reads that were kept against the reference genome of *A. percula* (GenBank Assembly ID GCA_003047355.1; Lehmann et al., 2018) using BWA-MEM V0.7.17 (Li and Durbin, 2009) and subsequently sorted, indexed, and filtered them according to mapping quality (>30) using SAMtools V1.8 (Li et al., 2009). Then, we assigned all the reads to read-groups using Picard Tools V2.20.7 (http://broadinstitute.github.io/picard/) and merged overlapping reads using ATLAS V0.9 (Link et al., 2017). We validated the mapping output using various statistics generated with BamTools V2.4.1 (Barnett et al., 2011). After mapping, we computed genotype likelihoods and estimated the major and minor alleles using the Maximum Likelihood Estimation method with ATLAS V0.9 (Link et al., 2017). We generated two datasets: one including all 89 individuals (including *A. akindynos* outgroup) and a second one comprising only the 67 *A. clarkii* individuals. We filtered the resulting two VCF files with VCFtools V0.1.15 (Danecek et al., 2011) to include only sites informative for all samples, with a minimum depth of 2 and a minimum Phred quality of 40.

### Population structure, admixture, and phylogenetic reconstruction

To explore the population structure and ancestral admixture of *A. clarkii*, we performed a principal component analysis (PCA) based on the covariance between individuals and calculated admixture proportions using PCANGSD V0.981 (Meisner and Albrechtsen, 2018). This method incorporates genotype likelihoods and connects the PCA results with the admixture proportions by detecting the most likely number of ancestral populations *K* based on the PCA eigenvectors. To further examine the genetic relationship between individuals, we built maximum likelihood phylogenetic trees for each chromosome with IQ-TREE V2.0.6 (Nguyen et al., 2015). We first converted the VCF files into multiple sequence alignment files (PHYLIP format) using vcf2phylip V2.0 (Ortiz, 2019). We then reconstructed each chromosome tree based on a GTR model and applied an ascertainment bias correction to account for the absence of constant sites in the SNP alignments. We assessed branch supports by performing ultrafast bootstrap approximation (UFBoot; Minh et al., 2013, Hoang et al., 2018) and SH-like approximate likelihood test (SH-aLRT; Guindon et al., 2010) with 1,000 replicates.

### Population genomics analysis in sliding windows and identification of outlier windows

To investigate the variation of genome-wide patterns of differentiation, divergence and diversity among *A. clarkii* populations, we estimated between-population differentiation (*F_ST_*), between-population absolute divergence (*d_xy_*) as well as population nucleotide diversity (*π*) in non-overlapping 5,000 SNPs windows using popgenWindows.py (https://github.com/simonhmartin/genomics_general). We summarised the variation across the 28 pairwise comparisons (*F_ST_* and *d_xy_*) and eight populations (*π*) using a PCA for each of the three statistics. We then normalised the resulting PC1 value calculated for each genomic window with a *Z*-transformation and used the *Z*-PC1 value as a single variable to describe genome-wide patterns of *F_ST_*, *d_xy_* and *π.* We identified outlier genomic regions by first applying a *Z*-transformation to the window-based *F_ST_*, *d_xy_* and *π* estimates to normalise the values across all populations and pairwise-population comparisons. We considered as outliers those windows with mean normalised values outside the 1^st^ and 99^th^ percentiles.

### Demographic modelling

We modelled the demographic history of *A. clarkii* using *fastsimcoal2* V2.6.0.3, a coalescent approach to infer demographic parameters based on site frequency spectrum (SFS; Excoffier et al., 2013). To simplify the models, we used the clusters identified with PCANGSD as populations. We grouped Indonesia, Philippines, and Taiwan into a single “Indo-Pacific Ocean” (IPO) population and Papua New Guinea and Solomon Islands into a “Pacific Ocean” (PAO) population. We considered Maldives and New Caledonia as two separate populations. We estimated the folded 2D SFS of the minor allele using whole-genome SNPs for each population using the vcf2sfs.py script (https://github.com/marqueda/SFS-scripts). We used the resulting 2D SFS to compare a total of eight distinct demographic scenarios (Figure 2). To first determine the origins of the four *A. clarkii* clusters (Maldives, IPO, PAO, and New Caledonia), we built four scenarios based on previous biogeographical reconstruction (Litsios et al., 2014). We modelled two scenarios with the PAO and IPO as the population of origin following a stepping stone process (models SS1 and SS2; Figure 2A and B), a scenario with the IPO population as the centre of origin (model CO; Figure 2C) and a scenario with a single ancestral population giving rise to both IPO and PAO populations, from which Maldives and New Caledonia respectively emerged (model SA; Figure 2D). We then used the best scenario explaining the origin of *A. clarkii* populations as the basis to build the four additional scenarios characterised by different timing of gene flow between populations: constant gene flow (Figure 2E), present-day exchanges between neighbouring populations (Figure 2F), gene flow after secondary contact (Figure 2G) and gene flow at the onset of the population splits (Figure 2H).

**Figure 2.**
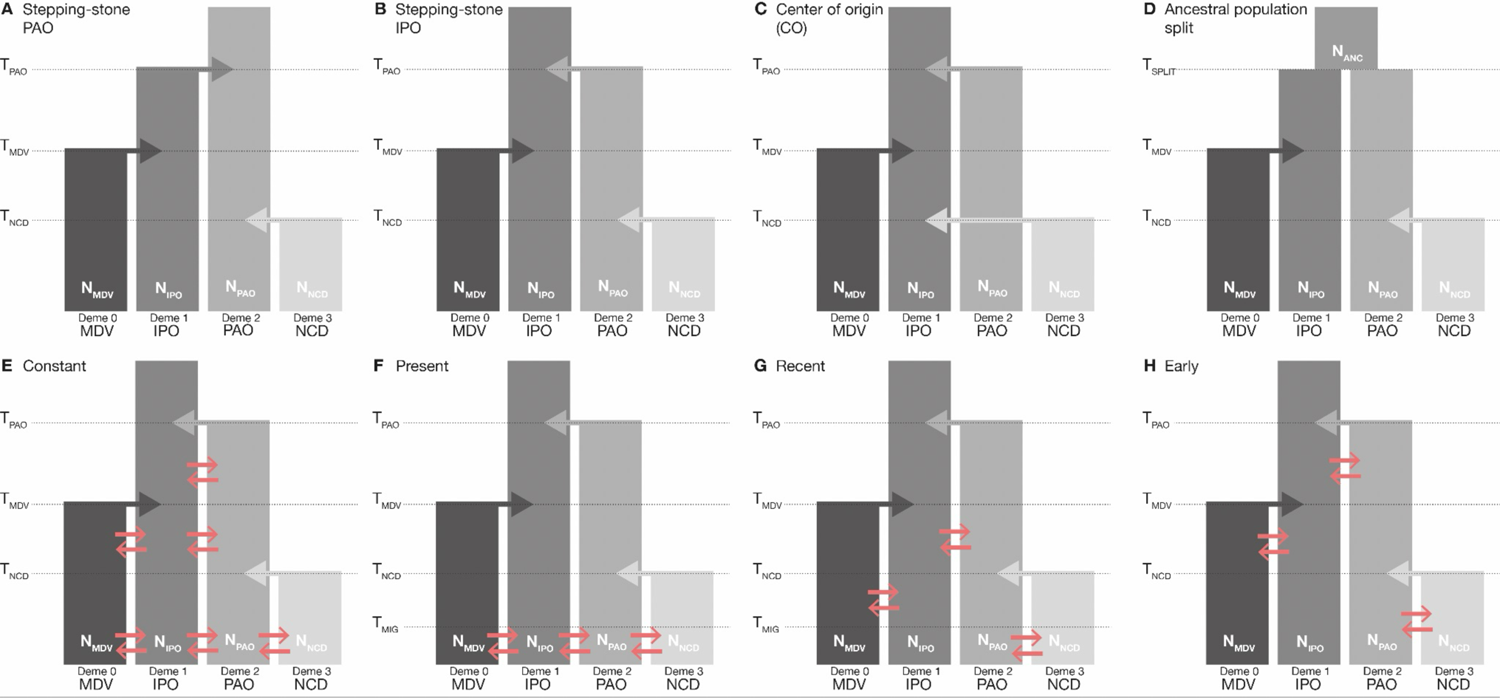
Demographic scenarios modelled with fastsimcoal26. Top scenarios depict the origins of *Amphiprion clarkii* populations. Stepping-stone models with either (A) the Pacific Ocean population as population of origin or (B) the Indo-Pacific Ocean population as origin; (C) Indo-Pacific Ocean population as centre of origin to all populations and (D) split from an ancestral population. Bottom models are characterised by different timing of gene flow. (E) constant gene flow; (F) present gene flow; (G) gene flow after secondary-contact (recent); (H) gene flow at the onset of the population split (early). Colonisation arrows (shades of grey) are depicted in coalescent time. The orange arrows represent migration (gene flow) between populations. IPO: Indo-Pacific Ocean (includes Indonesia, Philippines, and Taiwan); MDV: Maldives; NC: New Caledonia; PAO: Pacific Ocean (includes Papua New Guinea and Solomon Islands).

To estimate the demographic parameters (population split time, effective population size and migration rate), we ran *fastimcoal2* for each model using the observed multidimensional SFS. We specified an approximate mutation rate of 4E-8 (Delrieu-Trottin et al., 2017; Rolland et al, 2018) and a 5-year generation time (Buston and García, 2007). We performed 200,000 coalescent simulations (*-n*) to approximate the expected SFS, ran 50 cycles of the expectation-conditional maximisation (ECM; *-L*) to estimate the parameters and discarded entries with fewer than 10 SNPs per SFS count (-C). We ran each model 50 times to obtain the best combination of parameters estimates for our data. We selected the run with the highest likelihood and proceeded to 100 additional runs to obtain the likelihood distribution. We visualised the fit of the simulated to the observed SFS using the R script SFStools.r (https://github.com/marqueda/SFS-scripts/).

To select the best-fitting demographic model, we used the Akaike’s information criterion (AIC), which considers the number of parameters when comparing the likelihood of the different models. Since the AIC can lead to an overestimation of the most likely model if SNPs are not independent (not LD-pruned), we also compared the likelihood distribution of each model. Two models with an overlap of 50% or more in their likelihood range were considered as not differing significantly from each other.

Finally, for the best-fitting demographic model, we calculated the confidence interval of the parameters by running 100 replicates of parametric bootstrap. For each replicate, we simulated SFS from the maximum composite likelihood estimates and re-estimated parameters based on 30 independent runs with 200,000 dsimulations and 20 ECM cycles (Excoffier et al., 2013; Ortego et al., 2018). We calculated the 95 percentile confidence intervals of every parameter based on the highest likelihood run of each bootstrap replicate.

### Gene ontology

We investigated whether specific gene ontology (GO) terms were enriched in the genomic windows displaying outlier values of *F_ST_*, *d_xy_* and *π*, respectively. We used the 17,179 annotated genes inferred by Marcionetti *et al.,* (2019), among which 14,002 were annotated with *biological process* GO terms. We retrieved gene annotations for all the genes within (or overlapping) the outlier genomic windows and performed GO enrichment analysis by comparing those annotations to the complete set of protein-coding annotated genes of *A. percula*. We used the R package TOPGO V2.44 (Alexa and Rahnenfuhrer, 2020) to perform the gene ontology enrichment analysis based on Fisher’s exact test with the *weight01* algorithm. We defined a minimum node size of 10 and considered those GO terms with a raw *p*-value below 0.01 as significant, following the recommendations from the TOPGO manual.

### Image processing, colour proportions and redundancy analyses (RDA)

For individuals with available pictures (28 individuals from Indonesia, Maldives, and New Caledonia; see Table S1), we standardised the picture size (3,000 by 4,000 pixels) and resolution and corrected the white balance using Adobe Photoshop CC 2019. In addition, we set 28 morphological landmarks on each image using FIJI software (Schindelin et al., 2012). We used the landmarks to perform a Procrustes analysis using the R package MORPHO (Schlager 2017) and aligned the images to a mean shape, ensuring that each pixel represented the same body part in all images. Then, we removed the background masking the fish outline considering a spline regression along the landmarks outer body landmarks. We used those processed pictures in RGB format in all subsequent image analyses.

### Colour proportions

We summarised the variation in colour proportion by estimating the proportion of black, white, and orange across the body of each individual following Endler (2012). This approach considers which colours are adjacent to each other and classifies pixels into categories, based on colour transitions from one pixel to another. We classified the colour of each image into three categories (black, orange, and white) to estimate their proportions using a custom R script (see Data Accessibility).

### Redundancy analysis (RDA)

We used redundancy analysis (RDA; Legendre and Legendre 1998) to identify potential SNPs that could be associated with differences in colour pattern among populations based on 28 individuals that had both SNP and colour data available (Capblancq and Forester 2021). We first vectorized the images and concatenated the RGB channels to create a 36,000,000 x 28 matrix (i.e., 3,000 x 4,000 pixels, times the 3 channels and 28 individuals) and summarised the information in a PCA using R. Then, we used the first six PCA axes (hereafter referred as ColPCs) as the RDA explanatory variables, while constraining the axes with the population ID, the geographic coordinates of each population and the population genomic structure, which was quantified based on the loadings of the two first axes of a PCA on the genome-wide covariance among individuals. We performed RDAs on each chromosome separately, keeping only SNPs that were at least 1 kb distant from one another, and extracted the SNPs significantly correlated with at least one of the ColPCs. We estimated the Malahanobis distance of each SNP in the multidimensional RDA space and identified significant SNPs using the probability that a given SNP is an outlier compared to the normally distributed neutral SNPs in the RDA space based on a chi-squared test with an FDR correction (Benjamini and Hochberg, 1995). We randomly subsampled the same number of outlier SNPs across the genome and tested whether the geographic genomic structure was recovered using a limited number of SNPs. Finally, we looked for SNPs detected with the RDA analysis in the proximity (±10 kb) of pigmentation genes (Lorin et al., 2018) and summarised the SNPs information in a PCA, to compare the structure of the colour-related SNPs to the global population genomic structure.

## RESULTS

### Sequencing, mapping, and SNPs calling statistics

We obtained an average of approx. 108 million raw paired reads across samples, with the number of raw reads ranging from ca. 57 million to ca. 127 million (Table S2). After trimming low-quality regions and removing low-quality reads, each sample contained between 54 to 119 million paired reads, which corresponds to an estimated raw coverage of 6.4X to 20.2X (Table S2).

We mapped the reads to the reference genome of *A. percula* (Lehmann et al., 2018) with between 83% and 89% of reads mapping accurately (i.e., with pairs having the correct orientation and insert-size; Table S3). We further filtered the data by removing reads mapped with low-confidence and potentially redundant sequencing data arising from the overlap of paired reads to obtain a final coverage ranging between 4.5X and 14.4X (Table S3). Following these steps, we computed genotype likelihoods, estimated the major and minor alleles, and filtered the resulting VCF file to obtain a final number of 31,387,088 SNPs for the dataset including *A. clarkii* and *A. akindynos*, and 21,932,247 SNPs for the dataset containing only *A. clarkii* individuals.

### Population structure, differentiation, and admixture

We used a principal component analysis (PCA) on the covariance matrix between individuals to get an insight into the population structure of *A. clarkii*. We found a clear separation among the Pacific Ocean populations (Papua New Guinea, Solomon Islands and New Caledonia), the Indo-Pacific populations (Taiwan, Philippines, Indonesia) and Indian Ocean (Maldives) populations along the first axis, which explained 78.69% of the variance. Along the second axis, which explained 16.52% of the variance, we observed a split between the Indo-Pacific populations and the Indian Ocean population (Figure 1B), which suggests that the genomic structure coincides with the spatial distribution of *A. clarkii*.

Then, to investigate potential ancestral introgression among populations, we calculated the admixture proportion of each individual (Figure 1C). The optimal number of clusters inferred by PCAngsd was K = 5, respectively splitting populations from (1) the Maldives, (2) Indonesia (Tulamben and Manado) and Philippines, (3) Taiwan, (4) Papua New Guinea and the Solomon Islands and (5) New Caledonia. We observed substantially higher levels of admixture in the four Indo-Pacific populations compared to the Indian and Pacific Oceans populations, thus suggesting heterogeneous gene flow between *A. clarkii* populations across its geographic range.

### Genome-wide patterns of differentiation, divergence, and diversity

We investigated whole-genome patterns of differentiation, divergence, and diversity across the eight *A. clarkii* populations by calculating *F_ST_* and *d_xy_* between each pair of population as well as within-population nucleotide diversity (*π*) in 5,000 SNPs genomic windows (Tables S5-S7). We reported here the mean of these statistics across the genomic windows. We observed high disparities in mean *F_S T_* for every 28 pairwise comparisons, with estimates ranging from 0.003 (Indonesia-Manado *vs* Philippines) to 0.449 (Maldives *vs* Papua New Guinea; Figure 3A; Figure S1; Table S4). We found similar patterns for *d_xy_*, with mean values ranging from 0.042 (Papua New Guinea *vs* Solomon Islands) to 0.116 (Maldives *vs* Solomon Islands; Figure 3B; Figure S1; Table S4), while mean within-population diversity (*π*) ranged from 0.041 (Papua New Guinea) to 0.084 (Indonesia-Tulamben; Figure 3C; Figure S1; Table S4). To quantify the similarity of *F_ST_*, *d_xy_* and *π* genome-wide patterns between pairwise comparisons (for *F_ST_* and *d_xy_*) and populations (for *π*), we ran three PCA analyses using the pairwise comparisons (for *F_ST_* and *d_xy_*) or populations (for *π*) as variables, extracted the first principal component (PC1) and normalised it to summarise the pattern. We observed a high similarity in genome-wide variation for the three statistics across pairwise comparisons and populations (Figure S2). The first axes of each PCA explained a large proportion of variance (70.1% for *F_ST_,* 80.2% for *d_xy_* and 76% *π*). Mean *Z*-transformed values for each genomic window were highly correlated to their respective PC1 value (|r^2^| > 0.97), showing that PC1 is a good indicator of the windows-based score for each statistic. In summary, these results highlight heterogeneous differentiation, divergence, and diversity landscapes throughout the genome, but with comparable patterns between populations.

**Figure 3.**
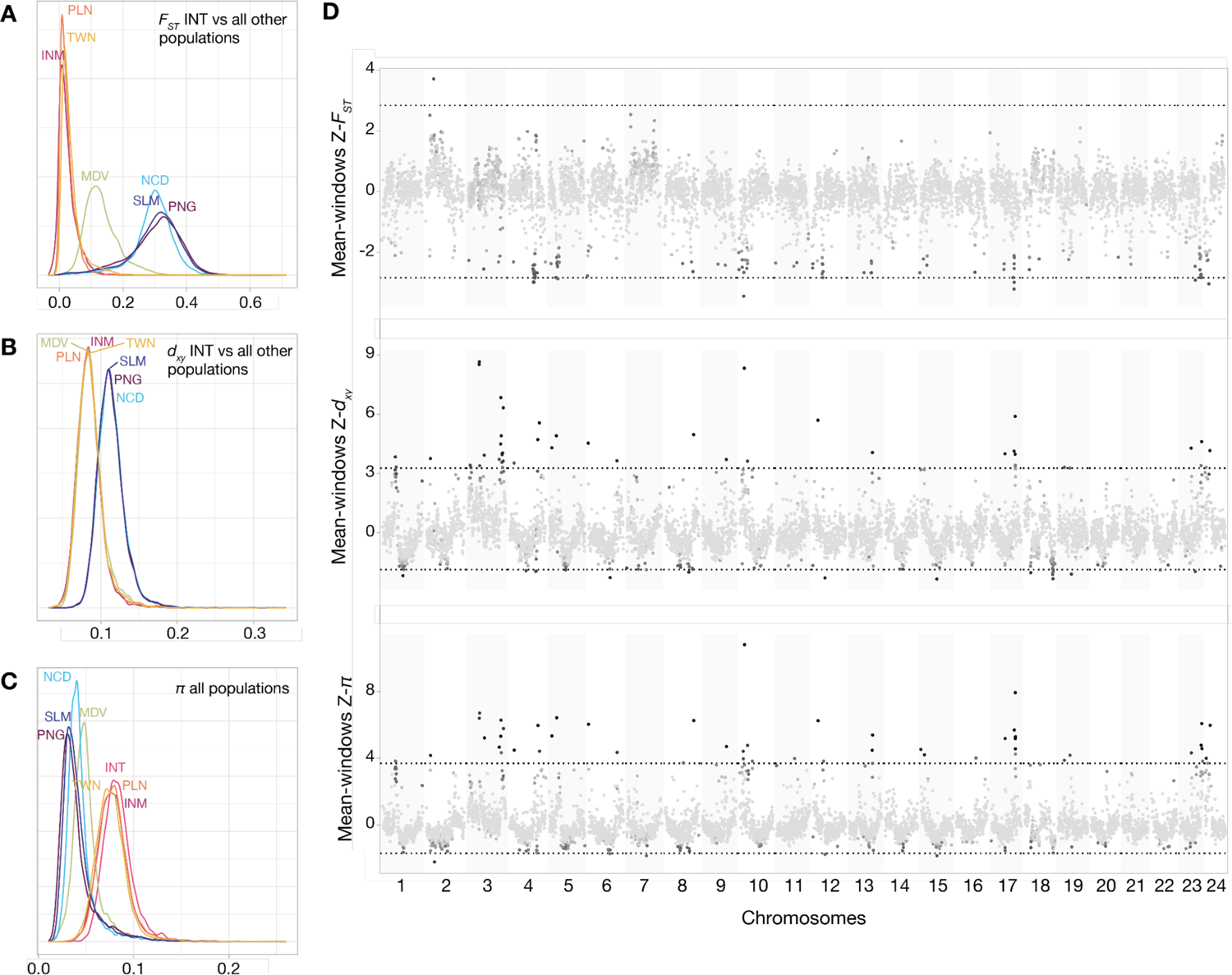
Genome-wide patterns of differentiation, divergence, and diversity. (A) Levels of differentiation (*F_ST_*), (B) divergence (*d_xy_*) and (C) diversity (*π*) within and among *Amphiprion. clarkii* populations calculated in 5,000 SNPs genomic windows. For clarity, we illustrated only pairwise comparisons with the Indonesian (INT) population. See Figure S1 for pairwise comparisons of *F_ST_* and *d_xy_* across all populations. (D) Mean Z-transformed estimates across pairwise comparisons (*F_ST_* and *d_xy_*) and populations (*π*) for each genomic window. Points are coloured according to the number of pairwise comparisons or populations displaying an outlier value (not included within the 1^st^ and 99^th^ percentiles, which are illustrated by the dotted lines). Black points mean that all 28 pairwise comparisons or all 8 populations show outlier values. Light grey points stand for no outlier value.

### Gene ontology of differentiation, divergence, and diversity outlier genomic windows

To identify genomic regions of potential importance in the evolutionary history of *A. clarkii*, we ran a gene ontology (GO) analysis on genomic windows with mean Z-transformed *F_ST_*, *d_xy_* and *π* found outside the 1^st^ and 99^th^ percentiles of values. We detected 11 outlier windows for *F_ST_*, 65 for *d_xy_* and 54 for *π.* Outlier windows were common to at least one-third of the pairwise comparisons or populations (Figure 3D; Table S8). We identified 117 genes in the *F_ST_* outlier windows (93 were functionally annotated), 542 in *d_xy_* (507 functionally annotated) and 553 in *π* (514 functionally annotated). We highlighted nine, ten and twelve significantly enriched terms (*p*-value < 0.01; Table S9) in the *F_ST_*, *d_xy_* and *π* outlier regions, respectively. The significant GO terms were associated with defence response to bacterium (GO:0042742) and lipoprotein metabolic process (GO:0042157) for both *d_xy_* and *π* outlier regions, while they were associated with proteolysis during cellular protein catabolic process (GO:0051603) and positive regulation of GTPase activity (GO:0043547) for *F_ST_* outlier regions.

### Demographic history analysis

To determine the origin of *A. clarkii* populations, we modelled four demographic scenarios characterised by different populations of origin and colonisation pathways (Figures 3A-D) using the coalescent modelling approach implemented in *fastsimcoal2*. The model with the highest likelihood was the stepping-stone model with an Indo-Pacific origin for the species (Figure 2B). This meant that the colonisation of both Maldives and Pacific Ocean populations came from the Indo-Pacific Ocean population followed by a more recent colonisation of New Caledonia from the Pacific Ocean population (see Table S10 for likelihood values).

Based on this model, we built four additional scenarios defined by different timing of gene flow (Figures 3E-H) to test for differential levels of migration between neighbouring populations. We obtained the highest likelihood for the model of constant gene flow (Figure 2E), which consisted of heterogeneous migration levels among neighbouring populations but constantly occurring through time (Figure 4). The three other scenarios (Figure 2F-H) had lower likelihood distributions that were not overlapping with the likelihood distribution of the best model (Figure 4B; Table S10) and were therefore rejected.

**Figure 4.**
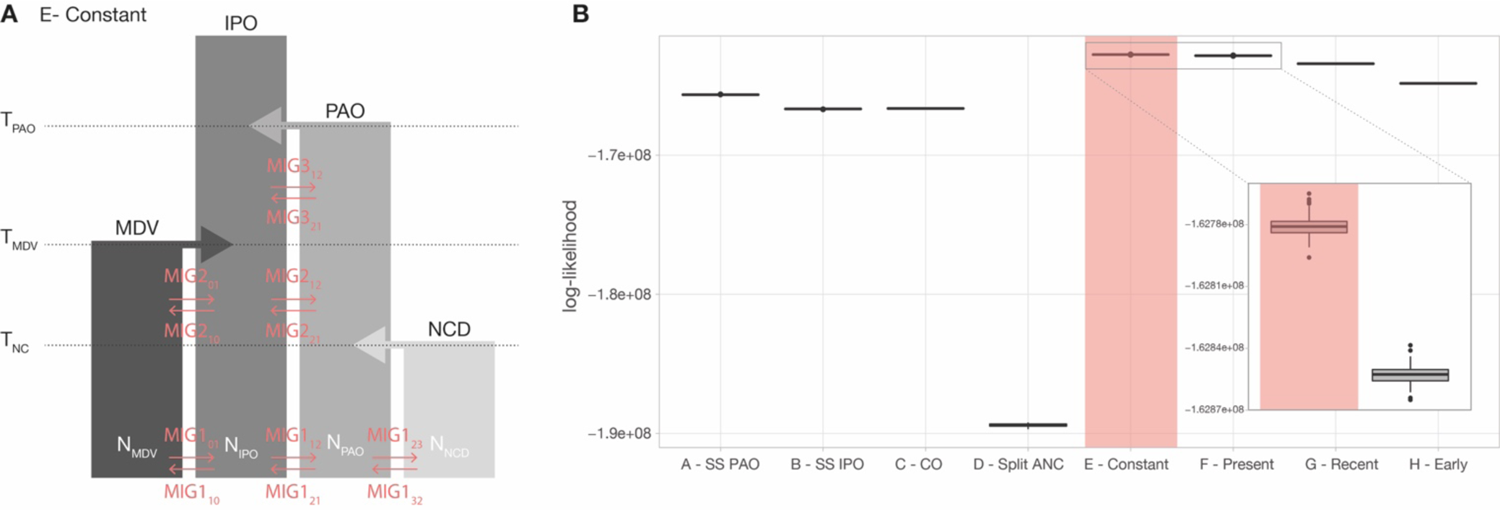
Demographic modelling results. (A) Scheme of the highest likelihood demographic scenario (E – Constant). Orange arrows represent gene flow between neighbouring populations. The parameter estimates for the best model are reported in Table 1. (B) Log-likelihood distributions for the eight tested demographic scenarios (see Figure 2). The best model is highlighted in orange, and a close-up of the two models with the highest likelihood distribution is displayed (E – constant, F – present). Likelihood distributions were obtained based on 100 expected SFS approximated using the parameters that maximise the likelihood for each model. Models with an overlap in their likelihood range indicate no significant difference between them. ANC: ancestral; CO: centre of origin; IPO: Indo-Pacific Ocean (includes Indonesia, Philippines, and Taiwan); MDV: Maldives; NC: New Caledonia; PAO: Pacific Ocean (includes Papua New Guinea and Solomon Islands); SS: stepping-stone.

**Table 1.** Parameter estimates for the best model with constant gene flow between neighbouring populations using the coalescent modelling approach implemented in *fastsimcoal2*.

| Population size |  | Parameter | Abbreviation | Estimate | 95% CI |
| --- | --- | --- | --- | --- | --- |
|  |  | Maldives | NMDV | 5,916 | [5,498; 8,549] |
|  |  | Indo-Pacific Ocean | NIPO | 19,858 | [19,082; 31,838] |
|  |  | Pacific Ocean | NPAO | 3,634 | [3,478; 5,282] |
|  |  | New Caledonia | NNCD | 3,849 | [3,617; 6,123] |
| Splitting time |  | New Caledonia | TNCD | 2,537 | [2,406; 4,397] |
|  |  | Maldives | TMDV | 11,591 | [11,591; 18,323] |
|  |  | Pacific Ocean | TPAO | 60,060 | [43,062; 524,780] |
| Migration rate per generation | After NCD split | MDV → IPO | MIG1_01 | 5.92E-05 | [3.98E-05; 6.40E-05] |
|  |  | IPO → MDV | MIG1_10 | 3.73E-05 | [2.37E-05; 4.51E-05] |
|  |  | IPO → PAO | MIG1_12 | 2.51E-05 | [1.27E-05; 2.51E-05] |
|  |  | PAO → IPO | MIG1_21 | 9.31E-05 | [6.50E-05; 1.02E-04] |
|  |  | PAO → NCD | MIG1_23 | 1.57E-04 | [1.15E-04; 1.79E-04] |
|  |  | NCD → PAO | MIG1_32 | 2.37E-04 | [1.53E-04; 2.80E-04] |
|  | After MDV split | MDV → IPO | MIG2_01 | 3.85E-05 | [1.68E-05; 5.14E-05] |
|  |  | IPO → MDV | MIG2_10 | 4.78E-05 | [2.78E-05; 5.88E-05] |
|  |  | IPO → PAO | MIG2_12 | 9.23E-08 | [3.92E-08; 1.69E-06] |
|  |  | PAO → IPO | MIG2_21 | 7.70E-05 | [4.51E-05; 7.70E-05] |
|  | After POA split | IPO → PAO | MIG3_12 | 9.86E-05 | [5.83E-05; 1.26E-04] |
|  |  | PAO → IPO | MIG3_21 | 1.16E-05 | [4.14E-08; 5.73E-05] |

We estimated the demographic parameters for the best model and highlighted that an initial split between Indo-Indian and Pacific populations (T_PAO_) occurred 60,060 generations ago. Subsequent splits of the Maldives (T_MDV_) and New Caledonia (T_NC_) populations occurred, respectively, 11,591 and 2,537 generations ago (Table 1). If we consider a generation time of five-years for clownfishes (Buston and García, 2007), our estimates corresponded to population divergence times of ca. 300 KYA (T_PAO_), 58 KYA (T_MDV_) and 12 KYA (T_NC_), respectively. We obtained the highest estimated effective population size for the Indo-Pacific Ocean population (N_IPO_), followed by the Maldives population (N_IPO_) and both Pacific Ocean (N_PAO_) and New Caledonia populations (N_NC_), the latter two having comparable sizes (Table 1). The estimated migration rates for the best model ranged between 9.23E-08 and 2.37E-04 (Table 1), with the highest migration rate corresponding to current gene flow between the New Caledonia and the Pacific Ocean populations (MIG2_32_). We obtained the lowest migration rate for the current gene flow between Indo-Pacific and Pacific Ocean populations (MIG2_12_; Figure 4A), although the confidence interval suggested a high level of uncertainty in the estimation of this parameter (Table 1).

### Coloration in *A. clarkii*

#### Colour proportions and patterns

We first outlined the coarse colour phenotype of *A. clarkii* individuals by calculating black, orange, and white proportions for each individual. We found a mean proportion for the black colour of 0.5, with values ranging from 0.19 (sample NC028) to 0.74 (sample GB030) between individuals. The orange proportion ranged from 0.04 (sample GB030) to 0.64 (sample NC028) with a mean of 0.31, and the white proportion displayed a mean of 0.19 with a minimum value of 0.08 (sample MV186) and a maximum value of 0.26 (MV033; Figure 5A, Table S12). The measurements confirmed the high variability in colouration patterns present in *A. clarkii* populations.

**Figure 5.**
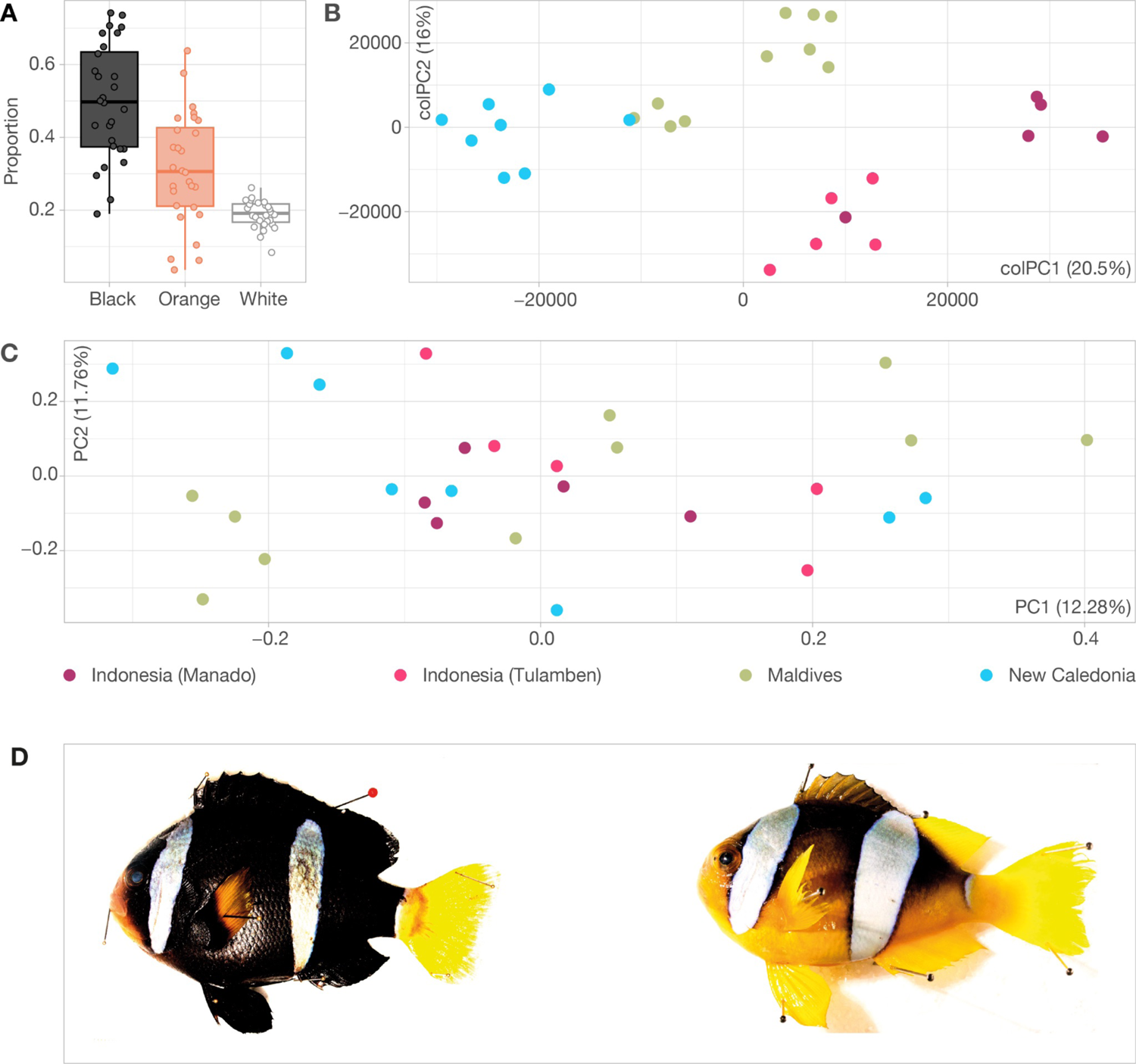
Colour proportions in *Amphiprion clarkii*. (A) Distribution of black, orange, and white colour proportions for the 28 individuals with available pictures (see Table S11). (B) Principal component analysis (PCA) summarising the colour of each pixel on two main axes for the 28 individuals. The first axis splits the different populations and explains 20.5% of the observed variation. The second axis explains 16% of the variation and splits the two Indonesian and Maldives populations. Coloured points represent the different populations. (C) PCA on the 224 significant SNPs highlighted with the RDA analysis showing that the population structure is lost. Coloured points represent the different populations. (D) Examples of individuals from Maldives (left) and New Caledonia (right) representing extreme phenotypes.

We further characterised the colour patterns of each individual based on the colour of each pixel and summarised the information in a PCA. The first six PCA axes on fish images explained more than 60% of the variance across images. The first axis (colPC1) explained 20.7% of the variance and differentiated yellow vs black colour variation of the ventral region, pectoral, pelvic, anal, and caudal fins as well as the extent and width of the white bands. It separated the populations of New Caledonia from Indonesian (INM and INT) populations. The second axis (colPC2) explained 16% of the variance and differentiated the orange vs black coloration of the ventral region, the intensity of yellow on the caudal fin and the shape of the middle white band. It separated the populations of Maldives from Indonesia – Tulamben (INT) populations (Figure 5B). The next four axis respectively explained 9.3%, 6%, 5.6% and 4.6% of the variance in coloration and differentiated the intensity of white on the middle band and the orange proportion of the ventral region (colPC3), the shape variation of the middle band (colPC4; Figure S3A), the colour variation on the dorsal and anal fins as well as the presence/absence of a peduncle white band (colPC5) and the dorso-ventral variation of the middle white band (colPC6; Figure S3B).

### RDA analysis

To identify genomic regions potentially involved in the difference in colour patterns among individuals and populations, we performed an RDA and removed the effect of neutral genomic structure and geographic distances. We detected 224 SNPs that had significantly larger Mahalanobis distances in the RDA space and could potentially be related to the colour pattern variation (hereafter referred to as “significant SNPs”). The average number of significant SNPs per chromosome was nine (corresponding to an average proportion of 9.7E-4; Figure S4). None of them were found in a genomic region related to pigmentation in other teleost fishes (Figure S5). The PCA performed on the 224 significant RDA SNPs shows a complete loss of population structure (Figure 5C). In contrast, a random sampling of 224 SNPs across the genome still recovered the expected genome-wide population structure (Figure S6).

## DISCUSSION

The extreme variability in multiple phenotypic and ecological traits makes *A. clarkii* an intriguing case among clownfish species. It spans the broadest geographic range of all clownfishes, can establish a mutualistic relationship with any of the ten host sea anemone species and displays a high level of colour polymorphism. We used population genomic, demographic and colour pattern analyses to assess the genetic cohesion between populations and to test different hypotheses about the colonisation pathways that enabled this species to reach its broad geographic range. Population genomic analyses of eight clownfish populations, revealed similar genomic landscapes and common genomic regions with a high level of differentiation and divergence across all populations, suggesting that common patterns of selection shaped their genomes. We further show that *A. clarkii* originated in the Indo-Pacific Ocean and spread to its current range using a stepping-stone mode of dispersal. We identified substantial gene flow throughout the demographic history of the species, despite being heterogeneous over time and among populations. Finally, colour patterns were distinct between two of the edge populations, but central populations displayed higher levels of colour variation that were associated with specific genetic variants. Our results suggest that populations of *A. clarkii* form a cohesive evolutionary entity and that the highly polymorphic patterns that characterise this species are not associated with genomic clusters of differentiation.

### Population structure, genomic landscape of differentiation, divergence, and nucleotide diversity

Our results based on whole-genome sequencing revealed substantial genetic structure among *A. clarkii* populations, with the highest level of differentiation between populations from the Western part of the geographic range (Maldives, Indonesia, Taiwan, and Philippines) and populations from the Eastern part of the distribution (Papua New Guinea, Solomon Islands and New Caledonia; Figures 1B, 2A, 2B; Table S4).

Populations from Indonesia, Taiwan and the Philippines displayed the highest nucleotide diversity (Figure 3), consistent with their central distribution providing them with further opportunities for connections to other populations. The New Caledonia population, which is on the edge of the distribution, displayed in contrast higher nucleotide diversity across the genome than both Papua New Guinea and Solomon Islands core populations (Figure 3C). A previous study on *A. clarkii* revealed that an edge population from Japan exhibited lower nucleotide diversity than the core populations (Clark et al., 2021), which was expected since populations from the edge of the geographic distribution usually face more substantial genetic drift and have a lower effective population size (*N_e_*), two factors known to decrease genetic diversity (Glémin et al., 2003; Kawecki, 2008). The higher nucleotide diversity that we found in the New Caledonian population might be a consequence of the constant gene-flow from neighbouring populations, as suggested in other coral reef fishes (Bay and Caley, 2011).

Mean differentiation (*F_ST_*) across windows was high among populations from different clusters compared to some other damselfish species at a similar spatial scale (Frédérich et al., 2014). This finding was expected because of the limited pelagic larval dispersal (Wellington and Victor, 1989; Ye et al., 2011), with larvae usually spreading within a perimeter of less than 27 km from the spawning site, even if long-distance migration has been observed previously (Pinsky et al., 2010). Furthermore, the social structure of clownfishes restricts larval settlement to not only a specific set of environmental variables but also to the presence of an appropriate sea anemone species. Even in the case of *A. clarkii* and its wide range of anemone hosts, this lifestyle further constrains the available settling sites and could reduce long-distance genetic exchanges between populations. Moreover, the observed population fragmentation might be a remnant of past sea-level fluctuations. The Pleistocene’s last glacial maxima (ca. 20,000 years ago) witnessed the formation of the Indo-Pacific barrier, a nearly uninterrupted land barrier (Fleminger, 1986) that induced a substantial reduction of the connectivity between the Indian and Pacific Oceans basins (Voris, 2000). The resulting habitat fragmentation potentially led to vicariance and subsequent genetic drift in the formerly separated populations. This mechanism potentially left behind a genetic signature in current populations, as observed in many fish species (Fitzpatrick et al., 2011; Winters et al., 2010) and invertebrates (Barber et al., 2000; Lessios et al., 2003), with few exceptions to that (Gaither et al., 2010; Horne et al., 2008). Compared to the high inter-cluster genetic structure, differentiation among populations from the same clusters was relatively low and comparable with other clownfish species (Dohna et al., 2018; Timm et al., 2012) and damselfishes (Frédérich et al., 2012; Liu et al., 2014). The limited differentiation across the Indo-Pacific cluster can be explained by the action of the North Equatorial Current (NEC), which splits into the northward-flowing Kuroshio current and the southward Mindanao current (Toole et al, 1990) and likely has homogenised genetic variation across populations, like other coral reef fishes from the Coral Triangle (Liu et al., 2019).

Differentiation (*F_ST_*) and divergence (*d_xy_*) were highly heterogeneous across the genome and generated similar genomic landscapes among populations (Figure 3D, S2). The heterogeneous distribution of functional elements and local recombination rate across the genome can influence the indirect selection strength and lead to highly heterogeneous patterns of *F_ST_* and *d_xy_* across the genome, as observed in various taxa (Burri et al., 2015; Pease and Hahn, 2013; Stankowski et al., 2019). Likewise, we observed high similarities of the genomic landscape across populations, which suggests that genomic characteristics were inherited from a common ancestor and thus maintained among the different populations (Vijay et al., 2017). Although no genomic window was considered as outlier across all 28 pairwise comparisons (Figure 3D) – suggesting that gene flow is homogenising the variance among populations – the detection of highly differentiated parts of the genome comparable across some populations could point to regions that are under selection and might be adaptive, potentially explaining the widespread distribution of *A. clarkii*. Interestingly, no outlier regions for the high levels of differentiation and divergence across populations were enriched for biological functions known to be related to colouration or preference for host sea anemone – two highly variable traits among *A. clarkii* populations. Although we cannot rule out that the observed peaks of differentiation could act as barrier loci (i.e., “islands of speciation”; Turner et al., 2005), it is more likely that they reflect various factors influencing the level of nucleotide diversity. Indeed, processes such as recombination rate, genetic drift and local adaptation are known to affect the nucleotide diversity, which in turn influences the level of differentiation (Cruickshank and Hahn, 2014; Hartasánchez et al., 2014; Noor and Bennett, 2009; Wolf and Ellegren, 2017).

### Demographic reconstruction of *A. clarkii* colonisation

#### Origins of A. clarkii populations

The demographic modelling based on SNPs data supported the hypothesis of a centre of origin of all *A. clarkii* populations in the Indo-Pacific Ocean and was favoured over a centre of origin located in the Papua New Guinea-Solomon Islands area (Figure 2, Table 1). According to this model, range expansion in *A. clarkii* was thus most likely achieved through a stepping stone process over multiple generations, like other clownfish species (Huyghe and Kochzius, 2018). Additionally, sea surface currents might have facilitated dispersal, as they are known to impact marine species connectivity (Kool, Paris, Barber, and Cowen, 2011). Expansion into new areas can thus be achieved even in species with characteristics linked to low dispersal ability. Indeed, a recent meta-analysis highlighted that life-history traits are not strong predictors of population structure and that traits such as pelagic larval duration were not good predictors of dispersal (Gandra et al., 2021). The migration pattern we uncovered for *A. clarkii*, is also consistent with the origin of the whole clownfish adaptive radiation, which started in the Indo-Pacific Ocean (Litsios et al., 2014; Santini and Polacco, 2006). The findings further support the possibility of range expansion and colonisation for species with limited dispersal ability. More generally, this pattern of colonisation coincides with the “Centre of Origin” hypothesis of the generation of the observed biodiversity (Briggs, 2003), which suggests that older lineages occur in the centre of the Coral Triangle and have longer evolutionary history compared to Pacific Ocean and Indian Ocean populations that were colonised more recently (Drew and Barber, 2009).

### Gene flow between A. clarkii populations

The best model supported the hypothesis of migration throughout *A. clarkii* demographic history (Figure 4). The low likelihood of the two models representing primary and secondary contacts of populations (Table S10) further supported that populations were constantly in contact with no interruption of gene flow between neighbouring populations. This result was somewhat unexpected since historical processes such as climatic oscillations across geological time are known to have significantly influenced the genetic structure of many marine species over time (Hewitt, 2000). Hence, we expected that gene flow in *A. clarkii* would have varied across geological time according to various environmental factors known to act as barriers in the marine environment, such as temperature, salinity, or oceanic currents (Oleksiak, 2019). Gene flow might thus have occurred shortly after populations split (Malinsky et al., 2015; Schumer et al., 2018) – or following a period without exchange between populations (secondary contact; Canino et al., 2010, Liu et al., 2012, Nikolic et al., 2020).

Although the migration level was heterogeneous across populations and time (Table 1, Table S11), the extent of gene flow was considerably higher compared to what was previously found between sympatric species of clownfishes (ranging from 1.29E-8 to 7.08E-7; Marcionetti 2021) and was more comparable to population-level gene flow in other marine species (ranging from 3E-6 to 4E-6; Nikolic et al., 2020). Additionally, *A. clarkii* populations diverged in the recent past, with at least 10 times lower divergence times among their populations than the divergence times found in sympatric clownfish species (about 45,697 generations; Marcionetti 2021). The prevalence of gene flow throughout *A. clarkii*’s demographic history and the comparisons of demographic parameters with other clownfish species suggest that populational processes – and not speciation processes – are driving *A. clarkii* diversification. Furthermore, the constant and substantial gene flow between *A. clarkii* populations builds on the pervasiveness of gene flow in the evolutionary history of clownfishes (Litsios and Salamin 2014, Gainsford et al., 2020, Schmid et al., 2022). The presence of past gene flow among the *A. clarkii* populations might have enabled the species to reach a high level of genetic variation by reassembling old genetic variations into new combinations (Marques, Meier, and Seehausen, 2019). The newly emerged variations might have enabled them to colonise various environmental niches and occupy all the ten host sea anemone species, similar to the ancestral hybridization events between clownfish species that have potentially triggered their adaptive radiation (Schmid et al., 2022).

### Colour proportions

*A. clarkii* individuals were more heterogeneous in orange and black colours than white colouration (Figure 5A). The colouration-based clustering highlighted that only populations at the extreme of the sampling distribution (i.e., New Caledonia and the Maldives) did not have overlapping colour patterns (Figure 5B). In contrast, the individuals from the populations at the centre of the distribution (Manado and Tulamben, Indonesia, i.e., Coral Triangle) displayed higher variability of colour patterns. The higher colour pattern diversity in central populations might be a consequence of their connectivity with populations from the Pacific and the Indian Oceans. Alternatively, it could be that the species of sea anemone host influences the resident clownfish species colouration, with darker clownfish found in sea anemones from the *Stychodactyla* genus (Militz et al., 2016). As such, the high richness of host sea anemones (Camp et al., 2016) in the Coral Triangle could explain the high variability in coloration of *A. clarkii*. Thus, the broad spectrum of orange and black proportions we found here, could be a consequence of the plasticity of those traits, as recently suggested in Indian *A. clarkii* populations (Devi et al., 2021). On the contrary, the narrow range in white colour proportion might result from strict genetic regulation of the trait, which we would expect since the white bands in clownfishes form at early developmental stages (Salis et al., 2019).

Substantial diversity in colouration was previously highlighted in coral reef species which underwent a burst of diversification, such as the Hamlets (*Hypoplectrus spp.;* Hench et al., 2022), and was also observed in other adaptively radiating species like the *Heliconius* butterflies (Edelman et al., 2019). Colour pattern is thus a crucial ecological trait under strong selection in various cases of adaptive radiations (Belleghem et al., 2017; Campagna et al., 2017; Stryjewski and Sorenson, 2017), and various factors such as environmental heterogeneity and context-dependent selection can have a substantial impact on the appearance of colour variations (Sirkia and Qvarnstrom, 2021). Further investigations are needed to evaluate the impact of colour patterns on the fitness of *A. clarkii* and its underlying genetic causes. This information will be crucial to understand the potential link between the colour polymorphism and the widespread geographic distribution of the species as well as its ability to interact with all the ten host sea anemone species.

### Conclusion: is *A. clarkii* a single species or not?

The high colour polymorphism displayed by *A. clarkii*, as well as its broad geographic distribution and ability to interact with the ten host sea anemone species, had been persistent arguments calling for further investigation of the phylogenetic status of the species (Litsios and Salamin, 2014). Although the contrasting colour patterns between the two populations at the extreme of the sampling distribution suggested that there is potential for ecological speciation, based on our whole-genome analyses, there is not enough evidence to consider *A. clarkii* as a complex of cryptic species. There are many ways to define what a species is (De Queiroz, 2007) and even more methodological approaches to delineate them (Petzold and Hassanin, 2020). A critical part of species delimitation is to consider morphological, behavioural, and ecological data in parallel to genomic data. We considered mainly genomic data in this study, and the incorporation of behavioural and ecological data is needed in the future. Indeed, field observations suggest that *A. clarkii* is an aggressive fish and is remarkably efficient at defending its host sea anemone against fishes feeding on its tentacles compared to other clownfishes. This bold behaviour might have enabled *A. clarkii* to colonise even smaller sea anemones neglected by other clownfish species (Santini and Polacco, 2006). Moreover, morphological data should be investigated more thoroughly. We did not consider more subtle variations in colour patterns, which might be a key feature that makes populations highly distinguishable since cryptic species often have divergent phenotypes that are hard to measure (Singhal et al., 2018). Another consideration for studies based on genomic data alone, is that in the past decade, various cases of speciation with gene flow have been found (e.g., Wang et al., 2016, Capblancq et al., 2019). This shows that in some cases, only a limited part of the genome with restricted gene flow may be necessary to form new species. Thus, special care should be taken when investigating gene flow to consider genes potentially critical for adaptation, i.e., that are under selection (Cadena and Zapata, 2021; Vaux, Bohn, Hyde, and O’Malley, 2021).

Since the results suggest that *A. clarkii* is a single species, this raises the question of the factors that enabled *A. clarkii* to reach such a broad distribution compared to all other clownfish species, despite their sedentary adult life and reduced larval dispersal time. This question is even more interesting considering previous suggestions of a trade-off between host specialisation (i.e., number of host sea anemone species) and environmental specialisation, with species interacting with a limited number of sea anemone species exhibiting a broader environmental niche (Litsios, Kostikova, and Salamin, 2014). However, the broad geographical range exhibited by *A. clarkii* might be a consequence of their ability to interact with the ten host sea anemone species or their colour polymorphism. If one (or both) of those traits enhanced dispersal, the subsequent mating of the high-dispersing individuals might reinforce the phenotype’s frequency and amplitude, leading to increased dispersal, a process known as spatial sorting (Comerford and Egan, 2022; Shine, Brown, and Phillips, 2011). This process can produce a shift in the phenotypes that improve dispersal against the potential negative impact on the organism’s fitness in the newly colonised habitat. This study thus opens new paths for understanding how polymorphism emerged in *A. clarkii*, and why – contrary to other clownfish species – it did not diversify into multiple more specialised species. Future work should include a more detailed assessment of the species’ ecology regarding environmental niche and interactions with the host sea anemone and other clownfish species.

## ACKNOWLEDGEMENTS

We would like to thank the Lausanne Genomic Technologies Facility for sequencing the samples and the DCSR infrastructure of the University of Lausanne for the computational resources. We also thank the staff at the Lizard Island Research Station for fieldwork support and acknowledge the Dingaal, Ngurrumungu and Thanhil peoples as traditional owners of the lands and waters of the Lizard Island region. Funding was from the University of Lausanne funds, Swiss National Science Foundation, Grant Number: 31003A-163428. FC was supported by an Australian Research Council (ARC) Discovery Early Career Research Award DE200100620 and Discovery Project DP180102363.

## DATA ACCESSIBILITY

Metadata were uploaded in GEOME (https://geome-db.org/workbench/project-overview?projectId=565) and the genomic data were deposited on NCBI (BioProject ID: PRJNA1025355) and will be made available upon acceptance of the manuscript. The scripts and pictures used for the colour analysis will be deposited on Dryad upon acceptance of the manuscript.

## AUTHORS CONTRIBUTION

MBS, NS and SS designed research. JB, FC, SH, FH, GL, AM, JO, CR and VT contributed to sample collection. MBS and SS performed research, lab work and analysed the data. AGJ carried out the coloration analyses. SS wrote the manuscript. All authors read, made corrections, and approved the final version of the manuscript.

## SUPPORTING INFORMATION

Appendix 1. Sampling permits information.

**Table S1. Information on sampled individuals.** Information about the samples used for this study. Only samples with available pictures were used for the color proportion analysis.

**Table S2. Sequencing and read processing statistics.** Number of paired reads (sum of reverse and forward reads) after sequencing and after quality filtering. We estimated the coverage by multiplying the number of raw reads with the read length, and divided it by the approximate reference genome size (900 Mb).

**Table S3. Mapping statistics.** Number and percentage of total reads and proper-paired reads mapping to Amphiprion percula reference genome (Lehman et al. 2018). Properly paired means that paired-reads (forward and reverse) orientation is correct and that the gap between them corresponds to the expected insert size (∼350 pb). The number of final mapped reads correpond to the number of reads after filtering and that were actually use for all subsequent analysis. We estimated the coverage by multiplying the number of final mapped reads with the read length, and divided it by the approximate reference genome size (900 Mb).

**Table S4.** Mean *F_ST_*, *d_xy_* and nucleotide diversity (*π*) for each pairwise population and population. Mean *π*, *F_ST_* and *d_xy_* across all 5000 SNPs genomic windows

**Table S5. Pairwise *F_ST_* for 5000 SNPs genomic windows.** Pairwise *F_ST_* for each genomic window across all chromosome (Chr). INT: Indonesia (Tulamben), INM: Indonesia (Manado), MDV: Maldives, NCD: New Caledonia, PNG: Papua New Guinea, PLN: Philippines, SLM: Solomon Islands, TWN: Taiwan, Out: Outgroup (*A. akindynos*)

**Table S6. Pairwise *d_xy_* for 5000 SNPs genomic windows.** Pairwise *d_xy_* for each genomic window across all chromosome (Chr). INT: Indonesia (Tulamben), INM: Indonesia (Manado), MDV: Maldives, NCD: New Caledonia, PNG: Papua New Guinea, PLN: Philippines, SLM: Solomon Islands, TWN: Taiwan, Out: Outgroup (*A. akindynos*).

**Table S7. Population nucleotidic diversity (*π*) for 5000 SNPs genomic windows.** Population nucleotidic diversity (*π*) for each genomic window across all chromosome (Chr).

**Table S8. Outlier genomic windows for *F_ST_*, *d_xy_* and nucleotidic diversity (*π*) value.** Outlier genomic windows for *F_ST_*, *d_xy_* and *π*. Start and end of each windows are listed. *Z*-Q01 and *Z*-Q99 correspond respectively to the quantile 1% and 99% for the Z-transformed (normalized) statistics. Outliers correspond to the number of pairwise population comparisons (for *F_ST_* and *d_xy_*) and populations (*π*) having outlier values for the given window. The mean Z-value is the mean across all pairwise population comparisons/populations of the *Z*-transformed statistics for the given window.

**Table S9. Gene ontology for nucleotide diversity (*π*), *F_ST_* and *d_xy_* outlier windows.** Significant gene ontology (GO) terms based on the weight01 algorithm using a classic-Fischer test to calculate the p-value. All significant GO terms (*p* <0.01) are listed for each of the outlier genomic windows based on *F_ST_*, *d_xy_* or *π* statistics. Number of annotated genes across the whole-genome, number of significant genes, number of expected genes and classic-Fischer p-value are indicated for each GO term.

**Table S10. Likelihood and AIC of the 8 demographic models compared with fastsimcoal 2.** The model column corresponds to Figure 3. Models are ordered according to AIC value.

**Table S11. Effective migration rate for the most likely demographic model.** Effective migration rate and effective population size (*N_e_*) for the most likely demographic model (E - Constant) as infereed with fastsimcoal2. Effective migration rate was calculated as (2 x *N_e_* x MIG) and corresponds to the number of gene copies per generation. All other estimates for the most likely model are reported in Table 1.

**Table S12. Colour proportions.** Colour proportions used for the calculation of lambda index across genomic windows. We calculated the proportions following the approach described in Engler et al. 2012. Pictures were available only for a subsample of the initial dataset.

---

**Figure S1.** Genome-wide patterns of differentiation (*F_ST_*) and divergence (*d_xy_*) among *A. clarkii* populations calculated in 5,000 SNPs genomic windows. (A) Pairwise *F_ST_* comparison with the Indonesian (Manado, INM) population. (B) Pairwise *d_xy_* comparison with the Indonesian (Manado, INM) population. (C) Pairwise *F_ST_* comparison with the Maldives (MDV) population. (D) Pairwise *d_xy_* comparison with the Maldives (MDV) population. (E) Pairwise *F_ST_* comparison with the Philippines (PLN) population. (F) Pairwise *d_xy_* comparison with the Philippines (PLN) population. (G) Pairwise *F_ST_* comparison with the Papua New Guinea (PNG) population. (H) Pairwise *d_xy_* comparison with the Papua New Guinea (PNG) population. (I) Pairwise *F_ST_* comparison with the New Caledonia (NCD) population. (J) Pairwise *d_xy_* comparison with the New Caledonia (NCD) population. (K) Pairwise *F_ST_* comparison with the Solomon Islands (SLM) population. (L) Pairwise *d_xy_* comparison with the Solomon Islands (SLM) population.

**Figure S2.** Heterogeneous genomic landscape across chromosomes but comparable among populations. Plots of *Z*-PC1 for *F_ST_*, *d_xy_* and *π* in 5,000 SNPs windows. PC1 explains 70.1%, 80.2%, and 76% of the variation in *F_ST_*, *d_xy_* and *π*, respectively. We plotted the *Z*-transformed PC1 values such that above-average values have positive values and below-average values have negative values.

**Figure S3.** Principal component analysis (PCA) summarising the colour of each pixel on two main axes for the 28 individuals. (A) PCA displaying the third (colPC3) and fourth (colPC4) axis. (B). (A) PCA displaying the fifth (colPC5) and sixth (colPC6) axis.

**Figure S4.** Distribution of significant SNPs highlighted by the RDA analysis across chromosomes. Grey bars display the proportion of significant SNPs per chromosome. The proportion of SNPs was calculated by dividing the number of significant SNPs by the total number of SNPs in a specific chromosome. The size of the orange dots is proportion to the number of significant SNPs found in each chromosome. The dotted horizontal orange line corresponds to the proportion of significant SNPs averaged across all chromosomes.

**Figure S5.** Karyotype displaying the distribution of significant SNPs highlighted by the RDA analysis (orange triangles) and the pigmentation related genes in teleost fishes (purple dots; Lorin et al. 2018). No SNPs were found within a 10-kb regions of pigmentation genes.

**Figure S6.** PCA on 224 randomly sampled SNPs showing that the population structure is still recovered. Coloured points represent the different populations. Blue: New Caledonia; green: Maldives; pink: Indonesia (Tulamben); purple: Indonesia (Manado).

