## Appendix 1 for "Gene flow throughout the evolutionary history of a polymorphic and generalist clownfish"

### Supplementary material

#### Appendix 1 | Sampling permits

Samples from Australia were obtained by Fabio Cortesi research group. Permits were granted by the Queensland fisheries (180731), the Great Barrier Marine Park (GBRMPA; G17/38160.1) and the

University of Queensland, Animal Ethics (QBI/304/16).

Samples from Solomon Islands, Papua New Guinea and Lizard Island were obtained by Cynthia Riginos’ research group. Sampling in the Solomon Islands was via the Australian Government’s Pacific Strategy Assistance Program and with the assistance of the Roviana Conservation Foundation (Solomon Islands Government Ministry of Education and Human Resource Development and Ministry of Fisheries and Marine Resources research permit to S. Albert). Sampling in Papua New Guinea was in coordination with the National Research Institute, the Department of Foreign Affairs and Immigration (Research Visa: 10350008304) and the Department of Environment and Conservation (Permit to Export Wildlife: 011318). Sampling at Lizard Island was under the Great Barrier Reef Marine Park Authority and Queensland Parks and Wildlife Marine Parks (permits: G08/26733.1, G08/28114.1, G09/31678.1, G10/33597.1, G11/34452.1, G11/34640.1) and the Queensland Department of Primary Industries (QLD General Fisheries Permit: 118636, 150981).

Samples from Indonesia, Maldives and New Caledonia were obtained by Nicolas Salamin’s research group. Permits were granted by the Environment Direction of the South (N°60912-895-2017/JJC) and North Province (N°1291-2017/ARR/DENV) of New-Caledonia, the Ministry of Fisheries and Agriculture of Maldives (OTHR30-D/INDIV/2016/538) and the XXX

Samples from Taiwan were obtained by Filip Huyghe with the permit number 03/F/0390B.

Samples from Philippines were obtained by Jimmy O’Donnell. Permits were granted by the Bureau of Fisheries and Aquatic Resources (Commodity Clearance 2016-20091), the Palawan Council for Sustainable Development (PCSD, GP 2016-03, Wildlife Transport Permit no. 2016-05-000062-DMO – Calamianes) and the Fisheries inspection and quarantine clearance (Permit no. 009391).
