## Supplementary figures and images for "Gene flow throughout the evolutionary history of a polymorphic and generalist clownfish"

### Supplementary Figure S1

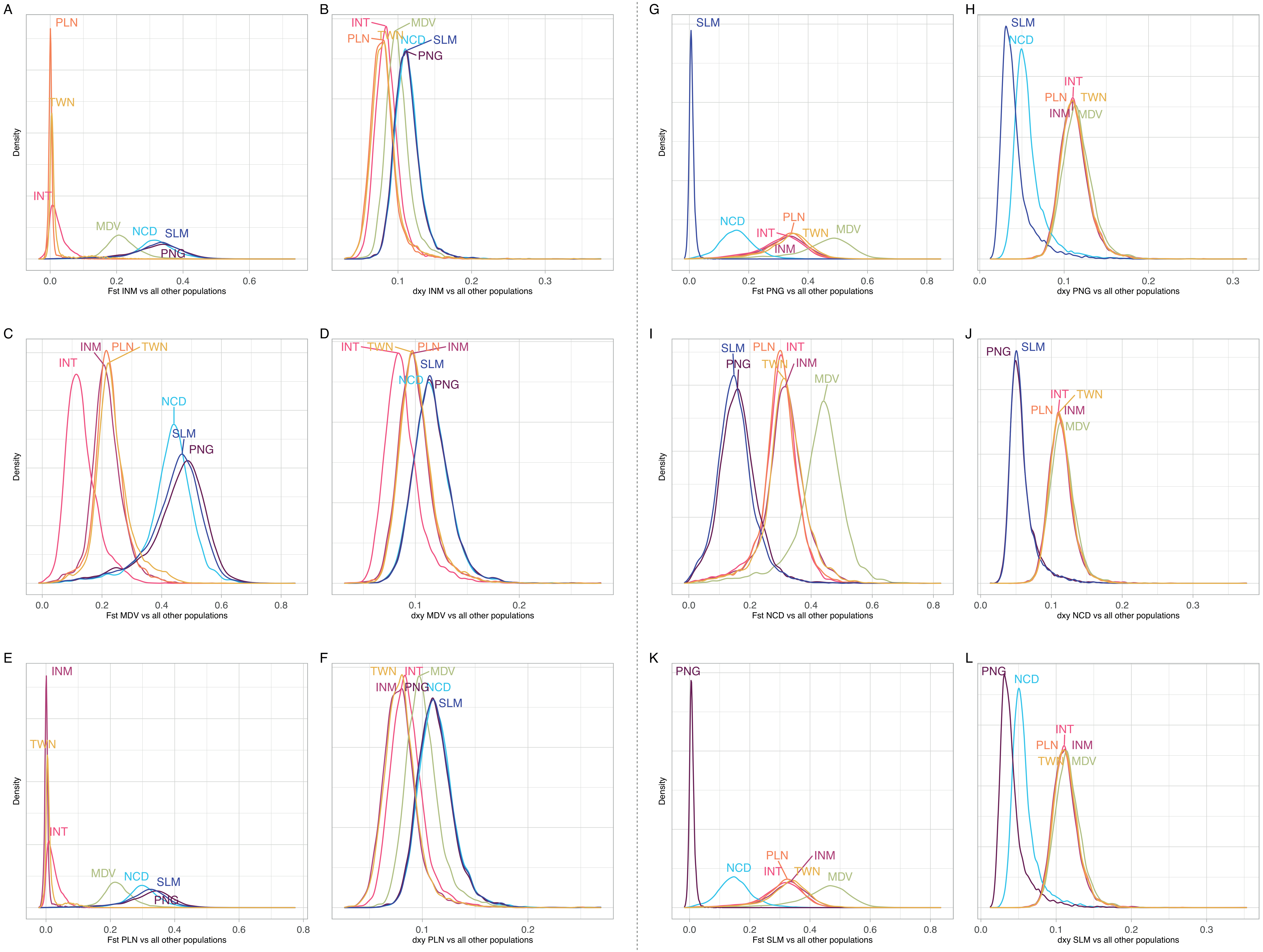

### Supplementary Figure S2

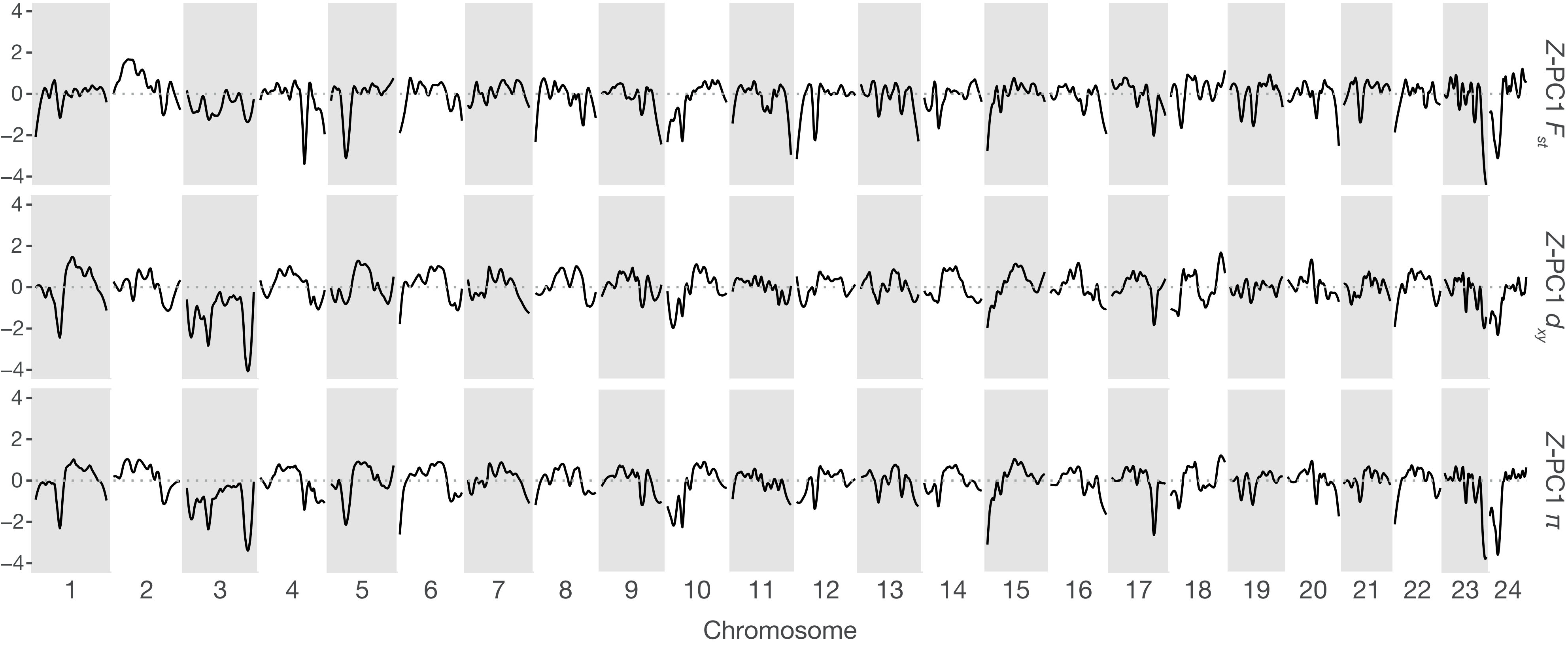

### Supplementary Figure S3

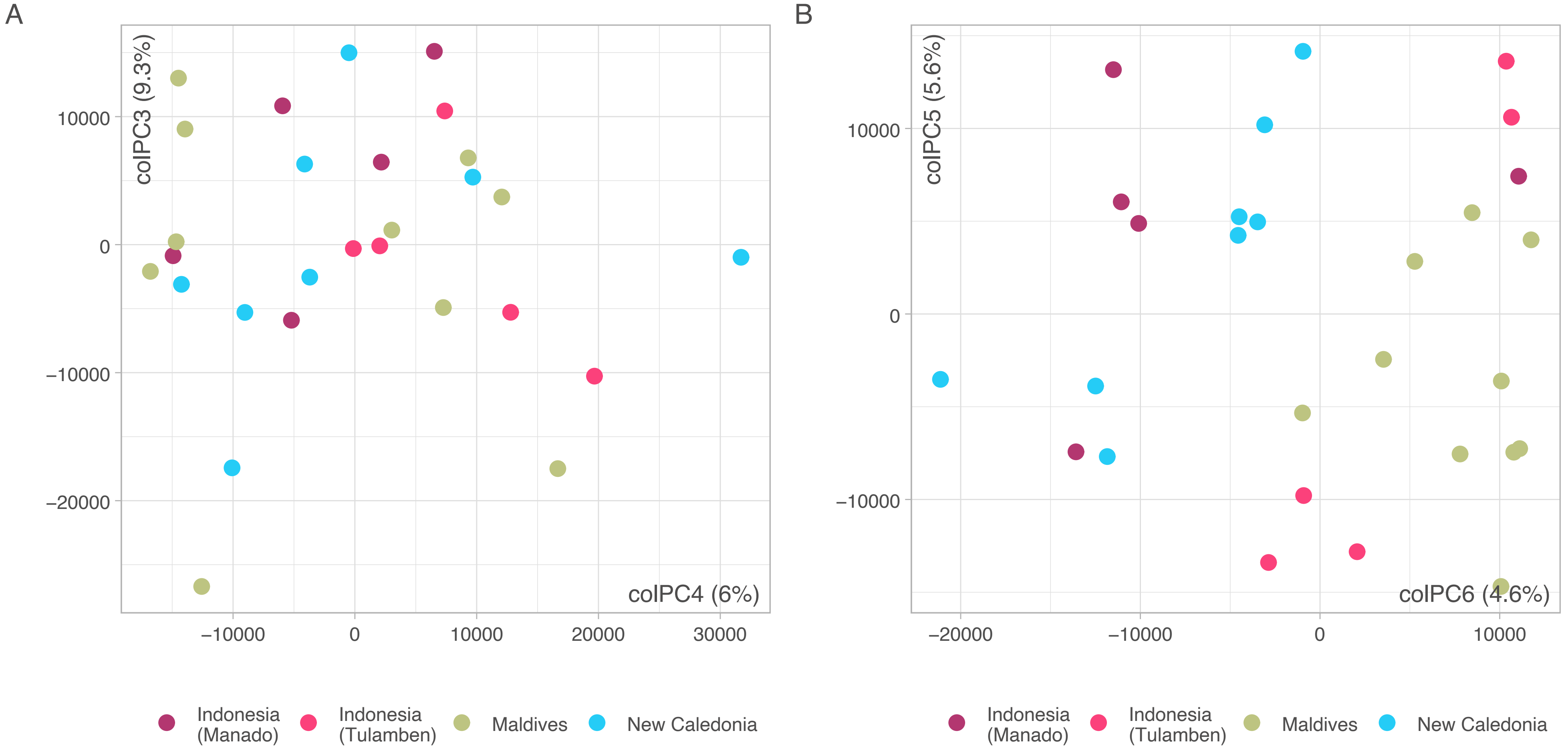

### Supplementary Figure S4

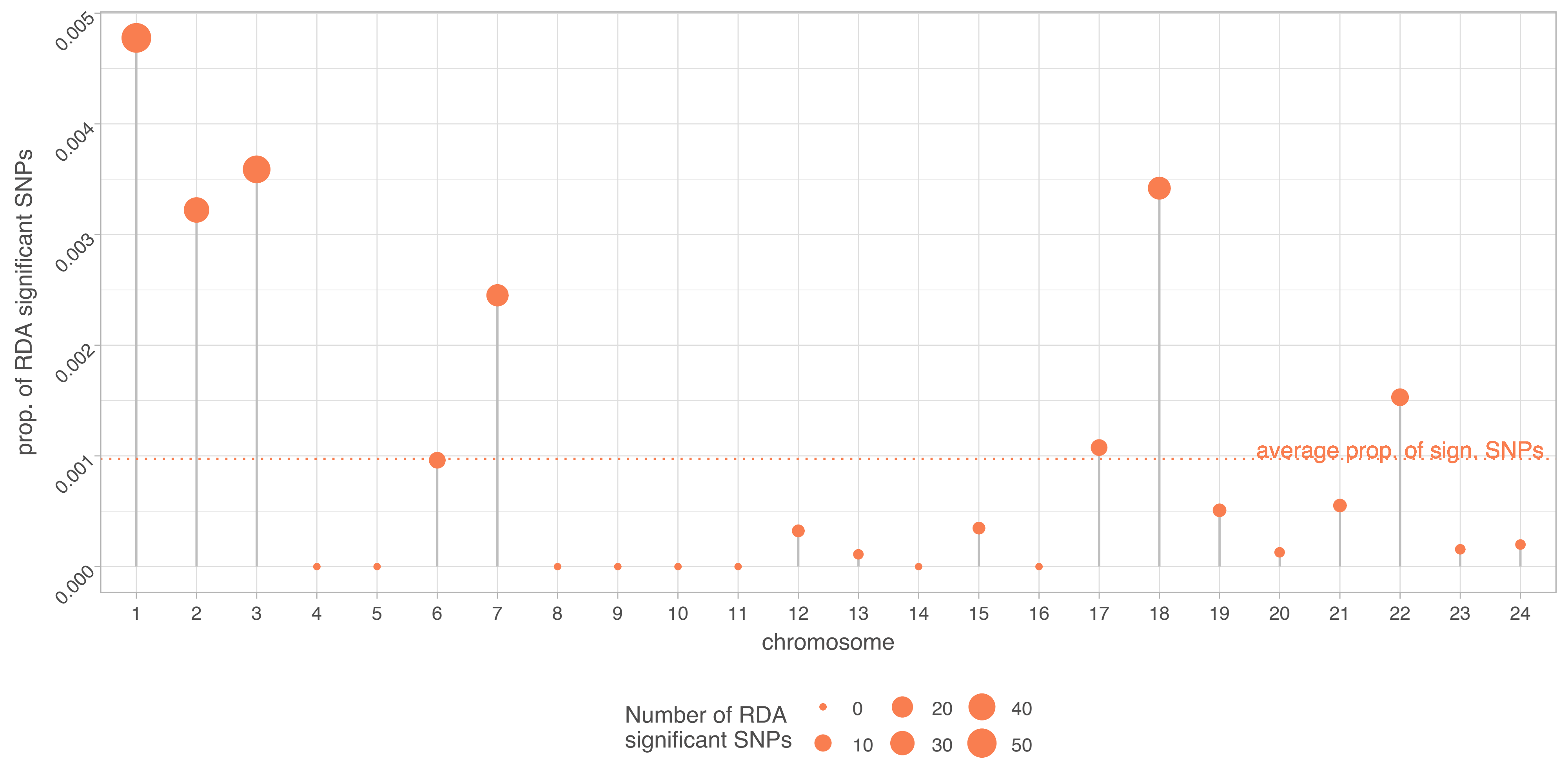

### Supplementary Figure S5

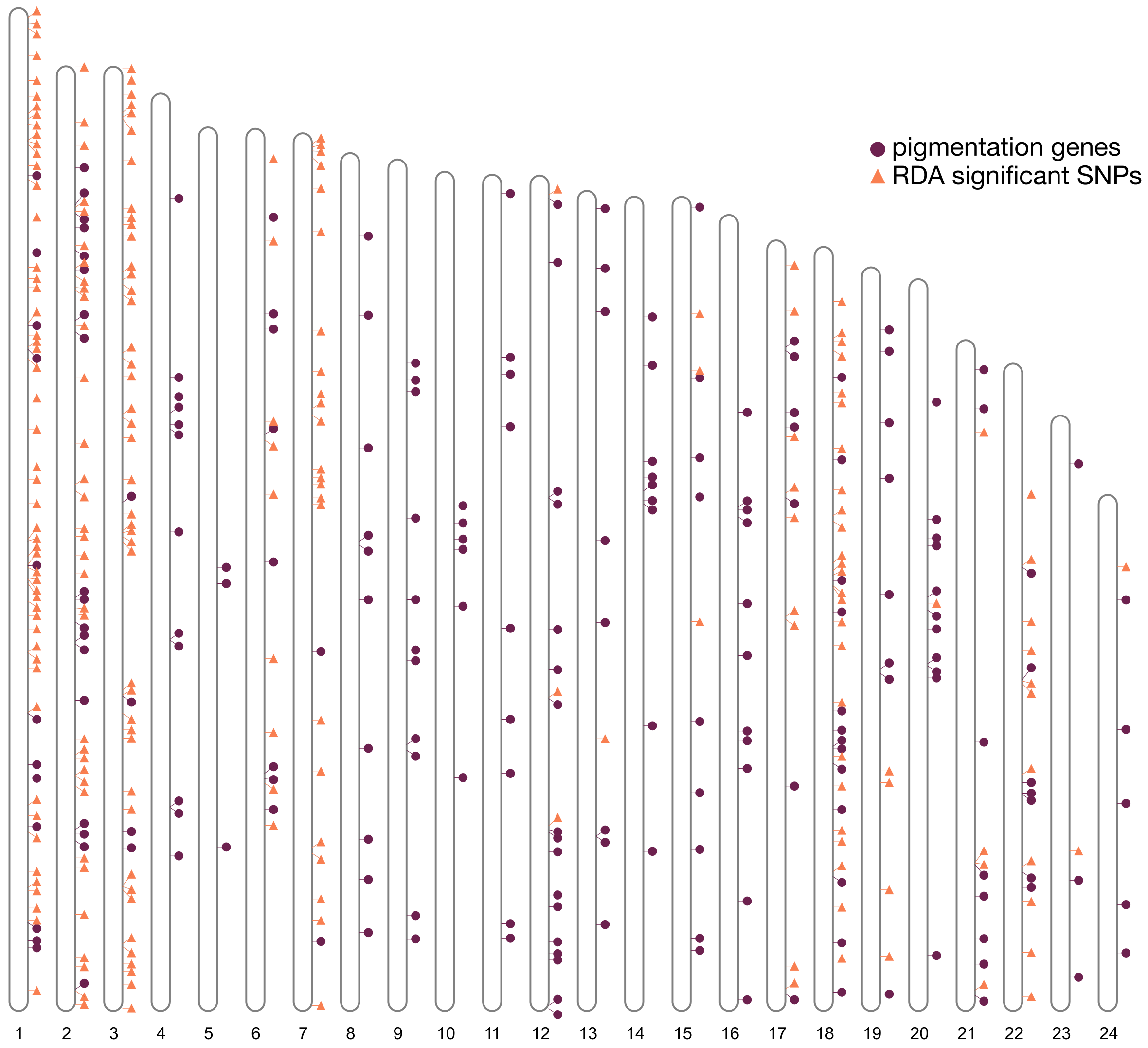

### Supplementary Figure S6

Randomely sampled SNPs (224)

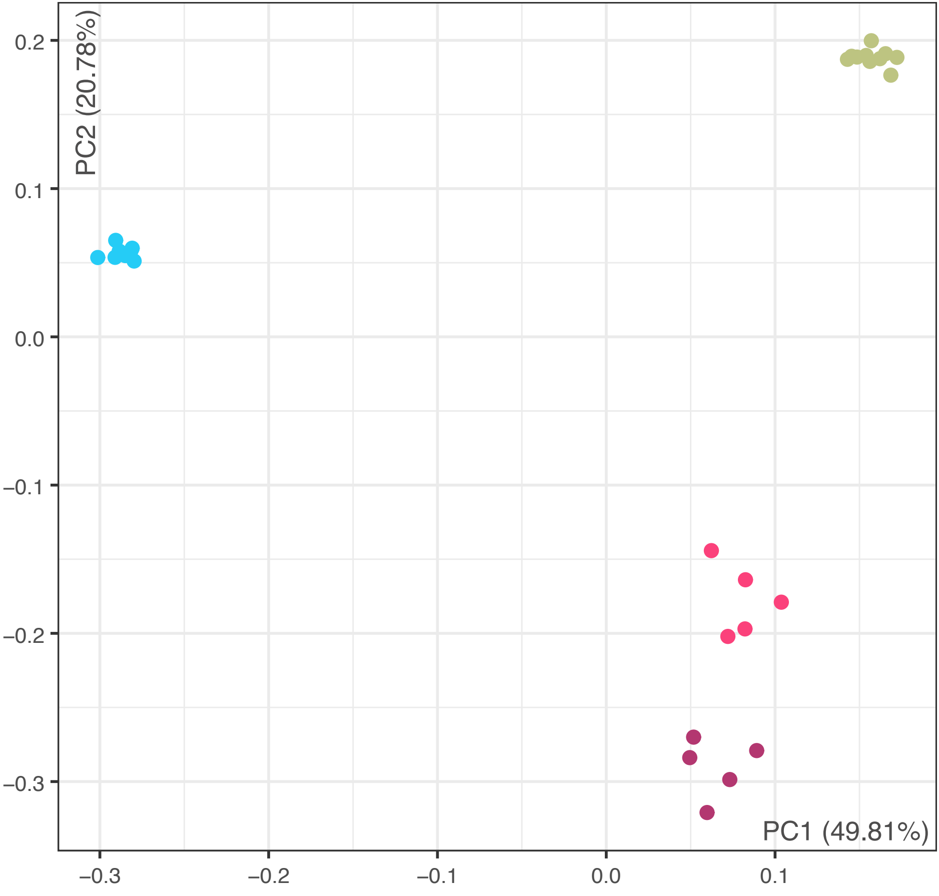
